# Comparative analysis of *Bacillus cereus* group isolates’ resistance using disk diffusion and broth microdilution and the correlation between resistance phenotypes and genotypes

**DOI:** 10.1101/2021.11.22.469644

**Authors:** Emma Mills, Erin Sullivan, Jasna Kovac

**Affiliations:** Department of Food Science, The Pennsylvania State University, The Pennsylvania State University, University Park, Pennsylvania, United States

**Keywords:** *Bacillus cereus* group, antimicrobial resistance, antimicrobial resistance genes, disk diffusion, broth microdilution, sensitivity, specificity

## Abstract

A collection of 85 *Bacillus cereus* group isolates were screened for phenotypic resistance to nine antibiotics using disk diffusion and broth microdilution. The broth microdilution antimicrobial results were interpreted using the CLSI M45 breakpoints for *Bacillus* spp. Due to the lack of *Bacillus* spp. disk diffusion breakpoints, the results obtained with the disk diffusion assay were interpreted using the CLSI M100 breakpoints for *Staphylococcus* spp. We identified significant (p < 0.05) discrepancies in resistance interpretation between the two methods for ampicillin, gentamicin, rifampicin, tetracycline, and trimethoprim/sulfamethoxazole. Antimicrobial resistance genes were detected using unassembled and assembled whole-genome sequences with Ariba and Abricate, respectively, to assess the sensitivity and specificity for predicting phenotypic resistance based on the presence of antimicrobial resistance genes. We found antimicrobial resistance gene presence to be a poor indicator for phenotypic resistance, calling for further investigation of mechanisms underlying antimicrobial resistance in the *B. cereus* group. Genes with poor sensitivity and/or specificity, as determined based on broth microdilution results included *rph* (rifampicin, 0%, 95%), *mph* genes (erythromycin, 0%, 96%), and all *van* genes (vancomycin, 100%, 35%). However, *Bc* (ampicillin, 64%, 100%) and *tet* genes (tetracycline, 67%, 100%) were highly specific, albeit moderately sensitive indicators of phenotypic resistance based on broth microdilution results. Only beta-lactam resistance genes (*Bc, BcII*, and *blaTEM*) were highly sensitive (94%) and specific (100%) markers of resistance to ceftriaxone based on the disk diffusion results, providing further evidence of these beta-lactams’ role in nonsusceptibility of *Bacillus cereus* group isolates to ceftriaxone.

**IMPORTANCE:** *Bacillus cereus* group includes human pathogens that can cause severe infections requiring antibiotic treatment. Screening of environmental and food isolates for antimicrobial resistance can provide insight into what antibiotics may be more effective therapeutic options based on the lower prevalence of resistance. Currently, interpretation of antimicrobial susceptibility testing results using the disk diffusion method is complicated by the fact that there are no standard disk diffusion resistance breakpoints defined for *Bacillus* spp. Hence, the breakpoints for *Staphylococcus* are often used in research studies. By comparing the results of disk diffusion interpreted using the *Staphylococcus* spp. breakpoints against broth microdilution interpreted using *Bacillus* spp. breakpoints, this study demonstrated that disk diffusion results interpretation with *Staphylococcus* spp. breakpoints are inconsistent. This study also provides new insight into the poor associations between antimicrobial resistance genotypes and phenotypes for the *B. cereus* group.

## INTRODUCTION

*Bacillus cereus* group, also known as *Bacillus cereus sensu lato* (*s*.*s*.), comprises spore-forming, Gram-positive rod-shaped bacteria commonly found in the environment and food (1–5). The *B. cereus* group is composed of eight genomospecies that are differentiated among each other based on the non-overlapping genome average nucleotide identity (ANI) of 95.2 (6). The *B. cereus* group genomospecies include *B. pseudomycoides, B. paramycoides, B. mosaicus, B. cereus sensu stricto* (*s*.*s*.), *B. toyonensis, B. mycoides, B. cytotoxicus, B. luti* (6). These eight genomospecies are phylogenetically separated into eight phylogenetic groups, defined based on the *panC* gene polymorphisms, that are consistent with the genomospecies taxonomy (7).

The *B. cereus* group species vary in their cytotoxicity towards human cells, which can be used as an indicator of pathogenicity. *Bacillus cereus* species (*Bacillus cereus s*.*s*.), which is known as a foodborne pathogen, has been reported to be overrepresented by strains with cytotoxic capability (8, 9). While the variation in cytotoxicity among *B. cereus* group phylogenetic groups has been documented, the differences in antibiotic nonsusceptibility among strains from different phylogenetic groups or genomospecies are largely unknown and are yet to be characterized (9, 10).

Scallan et al. (2011) estimated that 63,400 cases of *Bacillus cereus* foodborne illness occur in the United States annually (11). The prevalence of infections with *B. cereus* group bacteria may be underestimated due to the nature of foodborne illnesses and the passive surveillance of *B. cereus*. Although foodborne illness cases caused by *B. cereus* tend to be self-limiting, severe infections reported in the past required hospitalization and were lethal (12–15). For example, a 2003 outbreak linked to pasta salad contaminated with *B. cereus* consisting of 5 children led to one fatality (12). In 2008, another fatal case of *B. cereus* infection was reported due to consumption of contaminated pasta (10). *B. cereus* was also reported to cause other types of severe infections. In a retrospective study conducted at a French university hospital between 2008 - 2012, they found that nearly 30% of 57 patients hospitalized due to *B. cereus* infection had bacteremia and 28% had skin infection (16). Of the 57 hospitalized patients, 12% had died (16). Others have also reported *B. cereus* group infections associated with the hospital environment and supplies, including catheter-associated bloodstream infections in children in Japan and contaminated healthcare kits associated with cutaneous anthrax-like infections in newborns in India (17, 18). Although most *B. cereus* group infections may not require antimicrobial treatment, it is important to prepare for those that warrant critical care, particularly if novel infectious strains emerge within the *B. cereus* group.

The Centers for Disease Control and Prevention recommend ciprofloxacin, doxycycline, and beta-lactam antibiotics for treatment of inhalation exposure to *B. anthracis* (genomospecies *B. mosaicus* biovar Anthracis = *B*. Anthracis) (19). Severe non-anthrax *Bacillus cereus* infections are commonly treated with vancomycin, gentamicin, and clindamycin antibiotics (20–23). These antibiotics are widely used clinically due to the high prevalence of beta-lactamases found among *Bacillus cereus* group isolates (24–26), although specific beta-lactams (e.g., carbapenems) are also used for the treatment of clinical cases in combination (20, 27). Notably, there are reports of poor patient outcomes due to carbapenem resistance (14, 28). Recent studies have shown a high prevalence of resistance against penicillin, early cephalosporins, and 3rd and 4th generation cephalosporins in environmental isolates (2, 23, 29, 30). There have also been reports of a high prevalence of trimethoprim-sulfamethoxazole resistance in environmental isolates from Germany and China (23, 30). Understanding the prevalence of antimicrobial resistance among *B. cereus* group isolates can inform the selection of antibiotics for therapeutic use and potentially improve treatment outcomes.

In addition to understanding the prevalence of antimicrobial nonsusceptibility phenotypes, the information about underlying mechanisms of phenotypic nonsusceptibility is critical for the development of rapid diagnostic assays and for the surveillance of antimicrobial resistance. Studies on other foodborne pathogens, such as *Salmonella* and *Campylobacter*, have demonstrated the existence of a strong relationship between antimicrobial resistance genes (AMRg) or resistance mutations in housekeeping genes and phenotypic antibiotic resistance (31, 32). However, this relationship is yet to be characterized for the broad *Bacillus cereus* group, beyond *B*. Anthracis (19).

To study the correlation between individual AMRg and phenotypic nonsusceptibility, phenotypic susceptibility needs to be determined using standard interpretation breakpoints. Two commonly used methods for phenotypic antimicrobial susceptibility testing include disk diffusion and broth microdilution, which provide information on the zones of inhibition or minimum inhibitory concentration, respectively. Currently, the interpretation of the results of disk diffusion for *B. cereus* isolates is challenging due to the fact that current CLSI M45 guidelines only provide breakpoints for broth microdilution for *Bacillus* spp. (excluding *B. anthracis*). When utilizing the disk diffusion method, *Staphylococcus* spp. CLSI M100 breakpoints have often been applied for the interpretation of zones of inhibition for *B. cereus* group isolates (23, 30, 33). Furthermore, although broth microdilution clinical breakpoints for *Bacillus cereus* have been defined in the CLSI M45 for many antibiotics, there are still some antibiotics for which minimal inhibitory concentration (MIC) breakpoints are not available (21). It is unclear whether the broth microdilution results interpreted using *Bacillus* spp. breakpoints (CLSI M45) result in the same resistance interpretation as disk diffusion results interpreted using the *S. aureus* resistance breakpoints (CLSI M100). Hence, we sought to evaluate the consistency of resistance interpretation for 85 *B. cereus* isolates using disk diffusion interpreted using *S. aureus* breakpoints (CLSI M100) and broth microdilution results interpreted using *Bacillus* spp. breakpoints (CLSI M45). We further assessed the relationship between the detected antimicrobial resistance genes and antimicrobial resistance phenotypes to determine whether individual antimicrobial resistance genes are sensitive and specific markers of phenotypic resistance.

## RESULTS AND DISCUSSION

### The tested isolate collection was phylogenetically diverse and represented by all eight *B. cereus* group species

A total of 85 *Bacillus cereus* group isolates were available in our culture collection and were included in this study. These isolates had been previously collected from dairy-associated environments (65%), food processing environments (27%), and natural environments (7%) and are listed in Table S1. Whole-genome sequences for 85 isolates were obtained from the NCBI SRA database and were assembled (10, 34, 35). The average N50 of assembled genomes was 217 Kbp, the average genome size was 5.6 Mbp, and the average number of contigs longer than 1,000 bp was 129 contigs (Table S1).

Isolates were taxonomically identified using the average nucleotide identity (ANI) method implemented in the BTyper3 (36). Isolates belonged to 8 different species and the most abundant species were *B. mosaicus* (33%), *B. mycoides* (27%), and *B. cereus s*.*s*. (24%) (Table S1). Consistent with the species identification, most isolates were classified in *panC* phylogenetic groups VI *mycoides/paramycoides* (29%), II *mosaicus/luti* (26%), and IV *cereus sensu stricto* (26%) (Figure 1). There was a high MLST sequence type (ST) diversity among isolates, with 62 unique ST identified (Figure 1). Among these, 80% of identified STs were singletons. The most abundant STs were 1099, 1272, 222, and 1346, which belonged to phylogenetic groups IV *cereus s*.*s*., II *mosaicus/luti*, VI *mycoides/paramycoides*, and I *pseudomycoides*, respectively (Figure 1). The 5 isolates carrying Bt toxin-encoding genes (biovar Thuringiensis) were identified in three species, *B. cereus s*.*s*. biovar Thuringiensis (n = 2), *B. mycoides* biovar Thuringiensis (n = 2), and *B. mosaicus* biovar Thuringiensis (n = 1).

**FIGURE 1.**
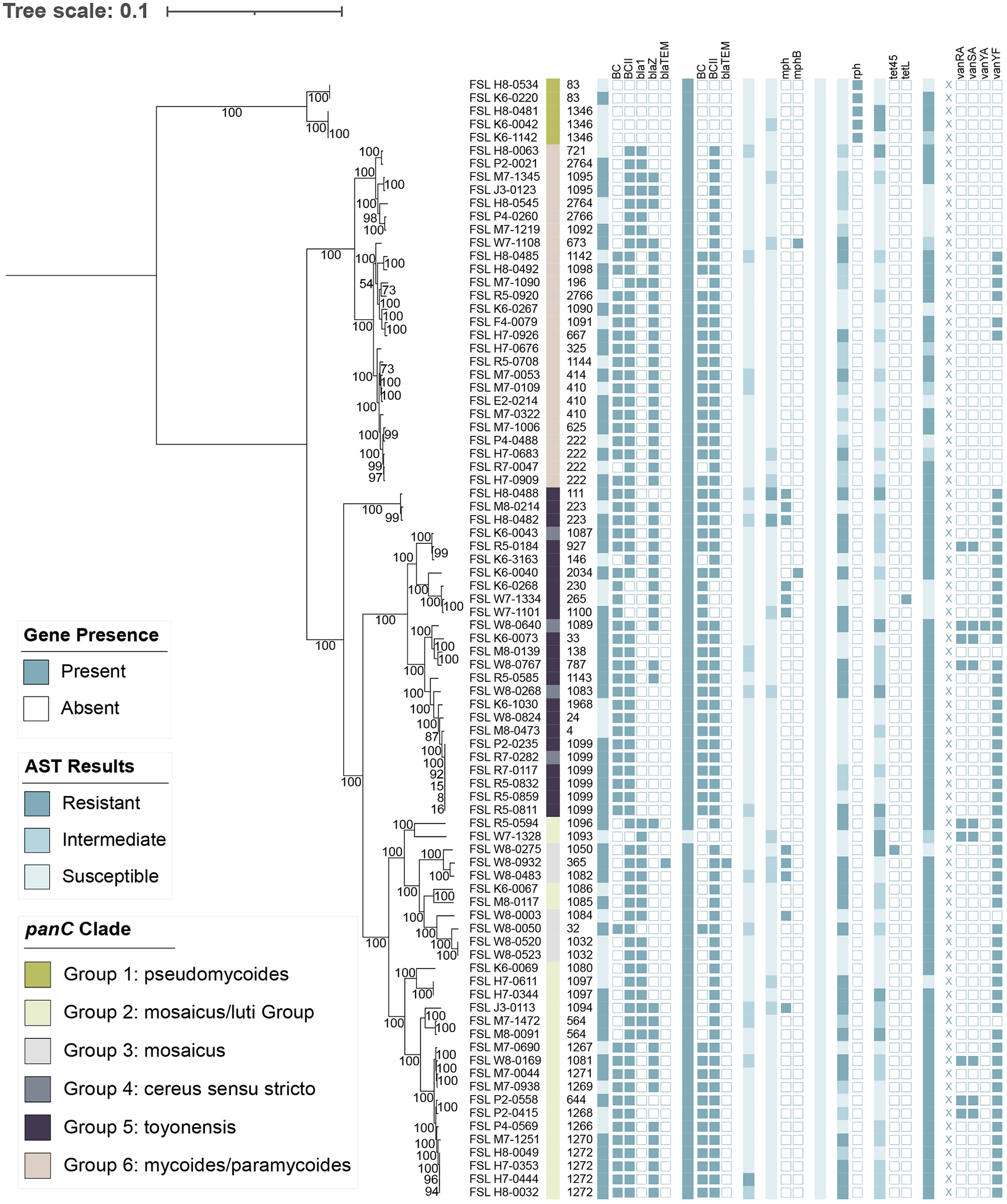
Phylogenetic tree for 85 *B. cereus* group isolates, annotated with the *panC* grouping (*panC*), MLST sequence type (ST), phenotypic resistance to ampicillin (AMP), ceftriaxone (CTX), ciprofloxacin (CPFX), erythromycin (ERY), gentamicin (GEN), tetracycline (TET), trimethoprim-sulfamethoxazole (TMP-SMX). The genes listed next to each antibiotic have previously been linked with phenotypic resistance to these antibiotics.

All isolates carried at least one gene from the *nheABC* operon, which is similar to reports from Ghana and China where they also reported a high prevalence of *nhe* enterotoxin genes in isolates obtained from dairy-associated environments and ready to eat foods, respectively (2, 37). In thirteen isolates’ genomes, we did not detect any *hbl* genes, whereas 65 isolates carried *hblABCD*, 6 carried *hblCD*, and only one carried *hblACD* (Table S1), alike abundance to previous works (2, 37). 40% of our isolates harbored cytotoxin K-2, nearly identical to the prevalence reported in Germany (37%) (30). The prevalence of these genes may be underreported due to the use of draft genomes for virulence gene detection.

### Abricate in conjunction with MEGARes 2.0 database resulted in the detection of the largest number of antimicrobial resistance genes

Antimicrobial resistance genes were detected using both unassembled reads with Ariba v2.14.6 and assembled genomes with Abricate v1.01., in conjunction with both MEGARes 2.0 and Resfinder databases. Abricate in conjunction with MEGARes 2.0 and Resfinder detected a median of 5 and 2 antimicrobial resistance-associated genes per isolate, respectively (Table S2). A search with Ariba in conjunction with MEGARes 2.0 and ResFinder databases resulted in a median of 2 and 0 detected antimicrobial resistance-associated genes per isolate, respectively. Given that the largest number of antimicrobial resistance-associated genes was identified using Abricate in conjunction with MEGARes 2.0 database, the genes identified by this program and database combination were used in further analysis (Table S2). The detection of acquired antimicrobial resistance genes by two different programs in conjunction with two different databases resulted in a different number of detected genes. Premkrishnan et al. also identified discrepancies in the number of detected antimicrobial resistance genes when using different methods for the detection of genes (38). This finding shows the importance of using multiple AMR gene detection tools to improve the detection of antimicrobial resistance genes (38).

### Beta-lactamase-encoding genes were detected in most isolates while other genes such as *tet, mph*, and *rph* were less prevalent

Nearly all isolates (93%) harbored genes *Bc* (n = 53) and/or a *BcII* (n = 76) that encode zinc Metallo-beta-lactamases. These genes were previously reported to confer resistance to penicillins, carbapenems and cephalosporins in *B. cereus* group species (25, 39, 40) (Table S2, Figure 1). Of the six isolates that did not have these genes detected, five belonged to *panC* phylogenetic group I (*pseudomycoides*) and ST-83 (n = 2), ST-1346 (n = 3), and ST-1093 (n = 1, *panC* group II) (Figure 1). These five group I (*pseudomycoides*) isolates uniquely carried an *rph* gene that was previously reported to confer resistance to rifampicin (41, 42) (Figure 1). Three beta-lactamase-encoding genes were identified among 85 tested isolates, including *bla1* (n = 25), *blaZ* (n = 48), and *blaTEM* (n = 1). These genes were detected in isolates from different *panC* groups and with different STs (Figure 1). All isolates carried a *fosB*, a thiol transferase-encoding gene that was previously associated with fosfomycin resistance in other bacterial species (43) (Table S2). Erythromycin resistance-conferring genes *mphB* (n = 2) and *mph* (n = 11) were identified predominantly in ST complexes ST-111 (n = 3), ST-23 (n = 3), and ST-365 (n = 2) (30) (Figure 1). Nearly all isolates belonging to *panC* group IV (*cereus sensu stricto*) carried the *sat* gene, predicting resistance to streptothricin, accounting for roughly half of the *sat* positive isolates (Table S2) (44). Only two isolates were genotypically predicted as resistant to tetracycline due to the presence of genes *tet45* and *tetL* (Table S1 and S2). These two isolates were identified as *B. moscaicus*, but belonged to different *panC* clades, III (*mosaicus*) and II (*mosaicus/luti*), respectively (Figure 1). Lastly, four genes associated with vancomycin resistance (45) were identified, *vanRA* (n = 9), *vanSA* (n = 9), *vanYA* (n = 1), and *vanYF* (n = 57) (Table S2). Genes *vanRA* and *vanSA* were detected in the same nine isolates; 5 group II (*mosaicus/luti*) and 4 group IV (*cereus sensu stricto*) isolates (Figure 1). The isolates positive for *van* genes were found in multiple *panC* groups. However, the 28 *van-*negative isolates were mostly found in *panC* group VI (*mycoides/paramycoides*) (71%) (Figure 1).

There have been few studies focused on characterizing the presence and abundance of acquired antimicrobial resistance genes (AMRg) in *B. cereus* group isolates (30, 46, 47). One study performed by Bianco et al. investigated the prevalence of AMRg in a collection of 17 *B. cereus* group, clinical blood isolates (46). Similar to our collection, there was a high prevalence of beta-lactamases in the Bianco et al. collection, specifically *Bla-1* and *Bla-2* (46). Our collection had a much higher percentage of the *blaZ* beta-lactam compared to the prevalence of this gene in the 17 clinical isolates (46). In our study, the Bianco et al. and Fielder et al study, *fosB* was identified in all isolates (30, 46). While Bianco et al. (46) identified a relatively high number of isolates carrying the specific *mphB* allele (erythromycin conferring), our collection showed a higher prevalence of the *mph* allele (13%) and a low prevalence of the *mph* allele (2%). Vancomycin resistance genes (*Gly-vanR-M, Gly-vanZF-Pp*, and *vanR-B*) were frequently represented among isolates reported by Bianco et al. (46) but were relatively rare among our isolates, with exception of *vanYF* (n = 57). Similarly to our findings, *tetL* was identified in only a singular clinical isolate in the previous study (46). Tetracycline resistance-conferring gene *tet45* was not identified in the clinical blood isolates from the Bianco et al. study (46), but was abundantly found in a study performed by Zhu et al. investigating virulence and resistance trends in *Bacillus cereus* probiotics (47). Zhu et al. (47) reported 8 of the 17 isolates carrying *tet45* and these isolates were identified as *B. cereus* (n = 7) and *B. thuringiensis* (n = 1). In comparison, we identified a singular isolate that carried *tet45* and this isolate was identified as *B. moscaicus*. Previous studies did not report *Bc, BcII, blaTEM*, and *rph* in isolates from the *B. cereus* group.

### Broth microdilution and disk diffusion often produced inconsistent results

Disk diffusion results for ceftriaxone and ampicillin were interpreted using EUCAST v11.0 breakpoints since the breakpoints for these antibiotics were not available in the CLSI M100-ED31:2021. The disk diffusion results for the remaining antibiotics (ciprofloxacin, erythromycin, gentamicin, rifampicin, tetracycline, trimethoprim/sulfamethoxazole, and vancomycin) were interpreted using the CLSI M100-ED31:2021 breakpoints. Although vancomycin zones of inhibition were measured and recorded (Figure 2A), breakpoints were unavailable in both EUCASTv 11.0 *Staphylococcus* spp. and CLSI M100-ED31:2021 *Staphylococcus* spp. guidelines.

**FIGURE 2.**
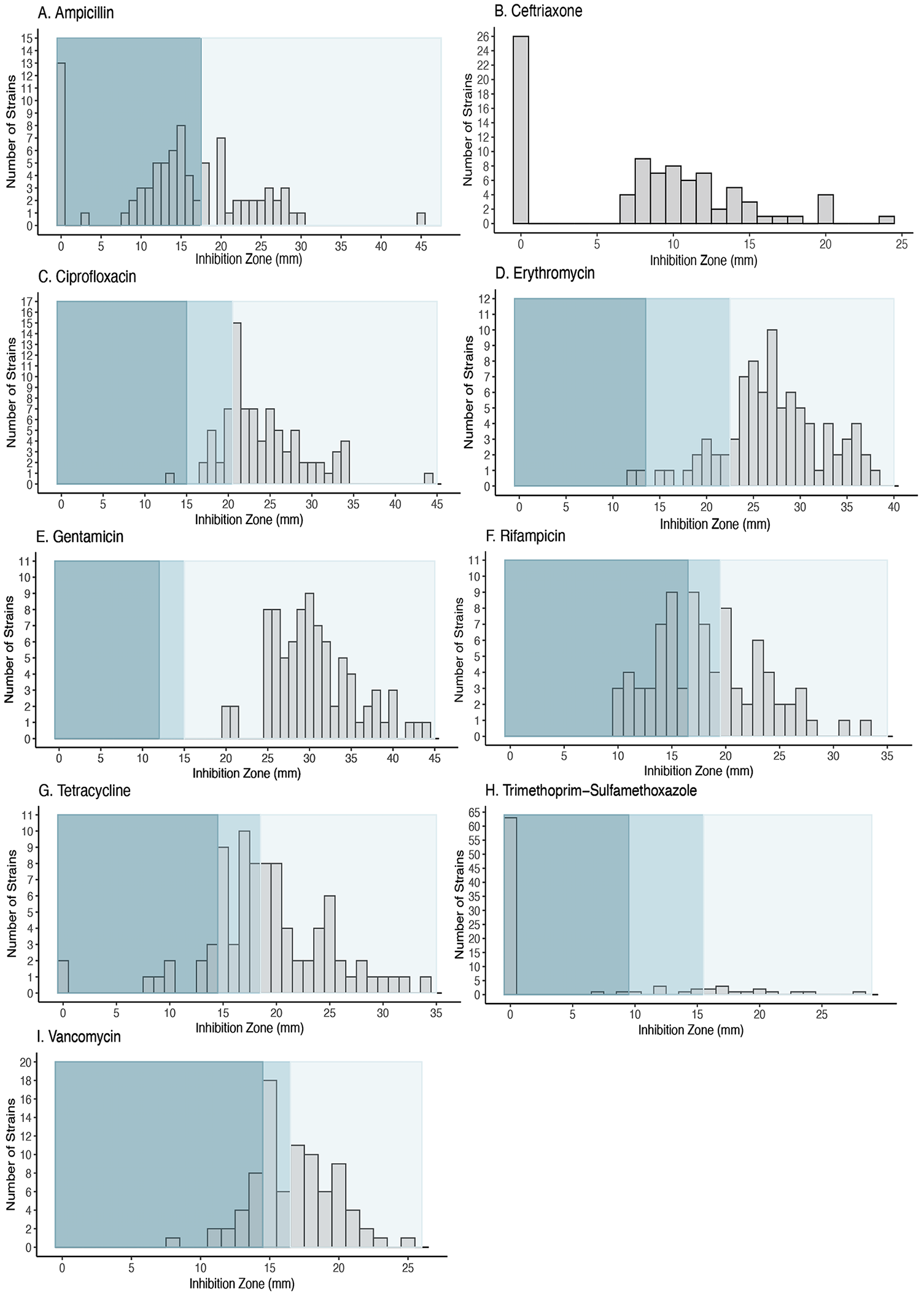
The distribution of isolates based on the zone of inhibition determined using disk diffusion for ampicillin (A), ceftriaxone (B), ciprofloxacin (C), erythromycin (D), gentamicin (E), rifampicin (F), tetracycline (G), trimethoprim-sulfamethoxazole (H), and vancomycin (V). The shades of grey-turquoise color show zones of inhibition that are considered resistant, intermediate, or susceptible based on the CLSI M100 or EUCAST v11.0 (ampicillin and ceftriaxone) disk diffusion breakpoints for *Staphylococcus* spp.

There was a high prevalence of beta-lactam resistance among tested isolates, with 62% of isolates resistant to ampicillin and 99% of isolates resistant to ceftriaxone (Table 1, Figure 1 and 2A). Only a few isolates were resistant to ciprofloxacin (1%, group II (*mosaicus/luti*)) and erythromycin (2%, group V (*toyonensis*)). All isolates were fully susceptible to gentamicin. 11% and 38% of isolates were resistant to tetracycline and rifampicin, respectively. The tetracycline and rifampicin-resistant isolates were sporadically found in different phylogenetic clades and among different sequence types (Figure 1). Lastly, nearly all isolates (78%) were resistant to trimethoprim-sulfamethoxazole.

**Table 1.**
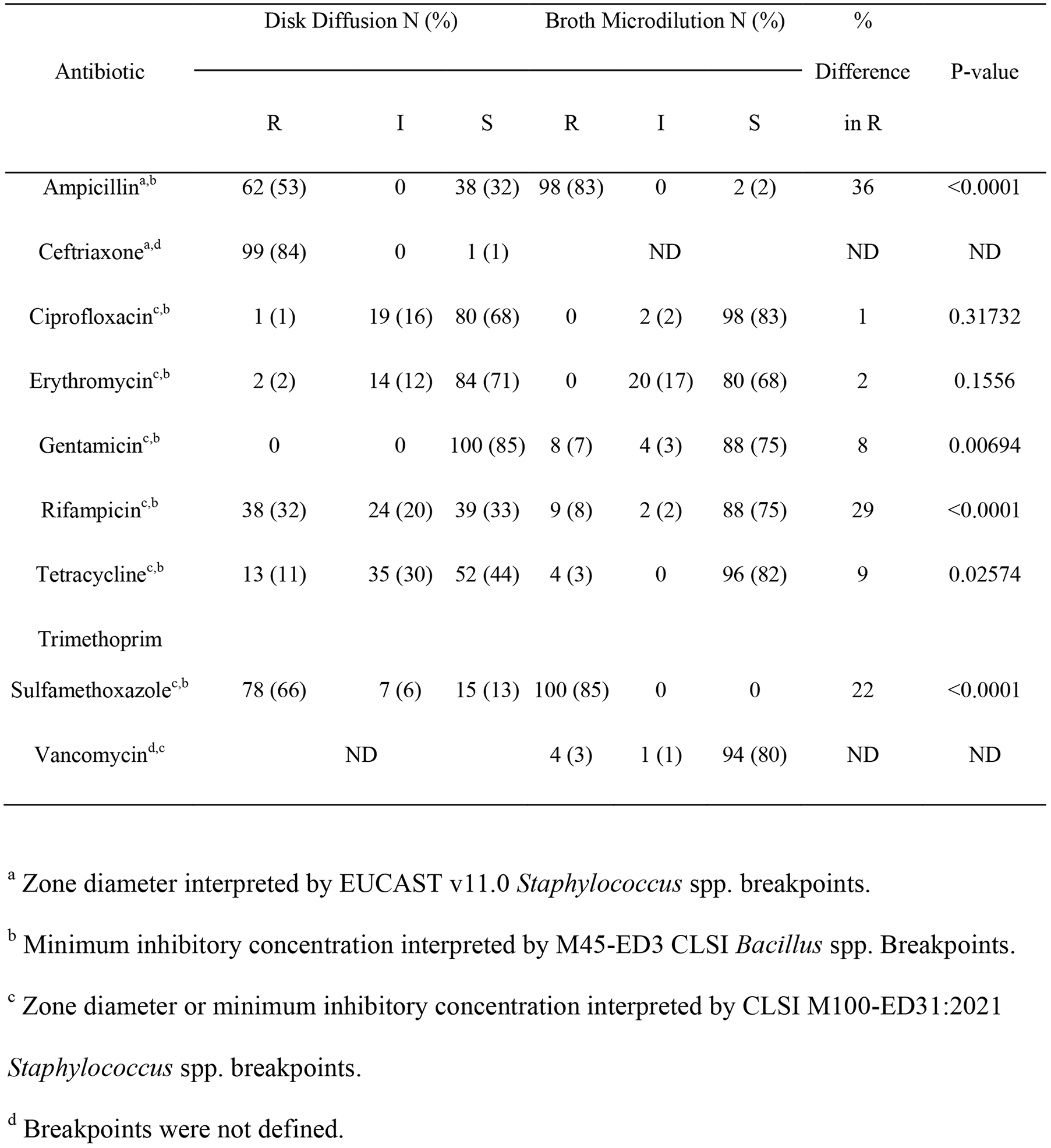
Antibiotic susceptibility of 85 *B. cereus* group isolates

Broth microdilution results for all antibiotics except vancomycin were interpreted using CLSI M45-ED3 *Bacillus* spp. breakpoints. Overall, there was a high prevalence of ampicillin (98%) and trimethoprim-sulfamethoxazole (100%) resistance (Table 1, Figure 3). No isolates were resistant to ciprofloxacin or erythromycin. There was a low prevalence of resistance to gentamicin (8%), rifampicin (9%), tetracycline (4%), and vancomycin (CLSI M100-ED31:2021 *Staphylococcus* spp.) among tested isolates.

**FIGURE 3.**
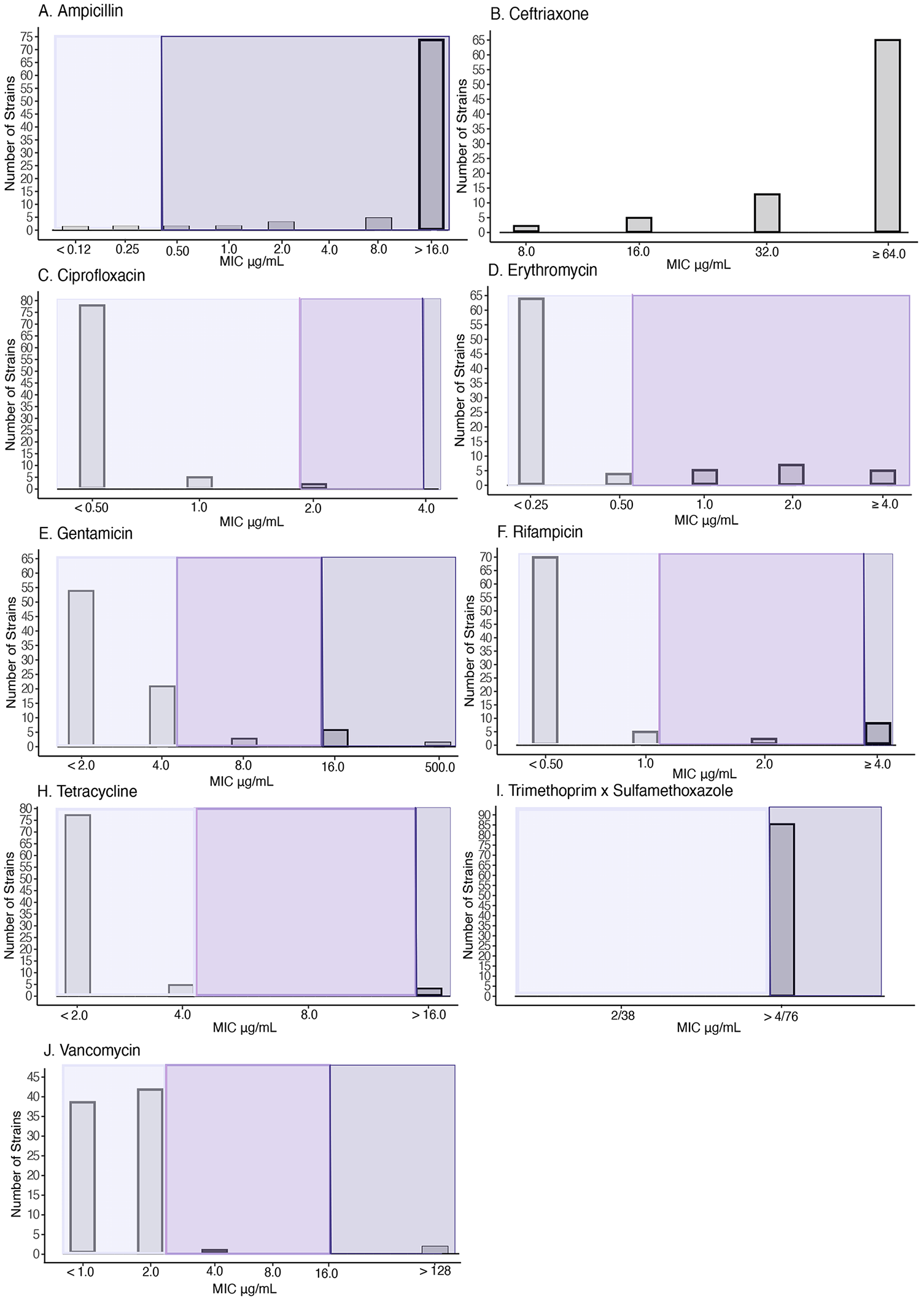
The distribution of isolates based on the minimum inhibitory concentration determined using broth microdilution for ampicillin (A), ceftriaxone (B), ciprofloxacin (C), erythromycin (D), gentamicin (E), rifampicin (F), tetracycline (G), trimethoprim-sulfamethoxazole (H), and vancomycin (V). The shades of grey-violet color show zones of inhibition that are considered resistant, intermediate, or susceptible based on the CLSI M45 broth microdilution breakpoints for *Bacillus* spp or CLSI M100 *Staphylococcus* spp. breakpoints (vancomycin).

When comparing the prevalence of resistance determined with the two AST methods, the prevalence of resistance to ampicillin (p < 0.0001), gentamicin (p = 0.00694), rifampicin (p < 0.0001), tetracycline (p = 0.02574), and trimethoprim-sulfamethoxazole (p < 0.0001) was significantly different when comparing disk diffusion and broth microdilution results (Table 1). There was statistically no difference in prevalence of resistance to ciprofloxacin (p = 0.31732) and erythromycin (p = 0.1556) when interpreted using the two methods. Since resistance to vancomycin and ceftriaxone were only measured by broth microdilution or disk diffusion, respectively, they were not included in the comparative analysis.

Our AST results obtained using disk diffusion were nearly identical to findings reported recently by Fiedler et al. that used similar disk diffusion methodology and EUCAST *Staphylococcus* v 6.0 breakpoints (30). In our study, as well as others, a low prevalence of resistance against ciprofloxacin, erythromycin, gentamicin, and tetracycline was reported (30, 37, 48). Further, previous studies also reported a high prevalence of resistance against third and fourth generation cephalosporins alike to our findings (30, 37, 48). Fiedler et al. reported a higher prevalence of resistance against ampicillin (99%), compared to the prevalence found in our study (63%) (30). In contrast, we identified a higher prevalence of trimethoprim-sulfamethoxazole resistance (78%), compared to Fiedler et al. (36%) (30).

The inconsistent prevalence of resistance between disk diffusion and broth microdilution for some antibiotics raises a question of the validity of the routinely applied *Staphylococcus* spp. breakpoints to *Bacillus cereus* disk diffusion results. The *Staphylococcus* spp. breakpoints have commonly been used for the interpretation of the resistance of *B. cereus* group isolates due to the lack of *Bacillus* spp. breakpoint availability (23, 30, 49). Although *Bacillus* spp. breakpoints have been defined by CLSI in M45-ED3 for broth microdilution; many studies have interpreted *B. cereus* group broth microdilution results using the CLSI M100. We did not identify any studies that had used *B. cereus* group isolates’ (excluding *B. anthracis*) broth microdilution results interpreted with *Bacillus* spp. breakpoints. Due to statistically significant differences in the prevalence of resistance when determined using the two AST methods, the application of *Staphylococcus* spp. breakpoints to interpret the resistance of *Bacillus* spp. isolates should be used with caution. Further studies need to be conducted to assess the validity of applying *Staphylococcus* spp. breakpoints to *B. cereus* group isolates.

### Comparison of antimicrobial phenotypes and genotypes revealed insufficient sensitivity and specificity for predicting resistance phenotypes based on the presence of resistance genes

Sensitivity and specificity were calculated for all genes identified by the Abricate/MEGARes 2.0 and the AST results for the corresponding antibiotics. The sensitivity and specificity were calculated for both disk diffusion and broth microdilution results to allow for comparison (Table 2). Among the ampicillin and ceftriaxone resistance genes, the gene encoding *BcII* (Metallo-beta-lactamase from a subclass B1) had the highest sensitivity for predicting ampicillin (96%) and ceftriaxone (90%) resistance, based on the disk diffusion results (Table 2). Sensitivity and specificity were not calculated for ceftriaxone using broth microdilution results due to the unavailability of breakpoints. Based on both the broth microdilution and disk diffusion results, the *BcII*-encoding gene had low specificity, 0% and 22% respectively, to ampicillin specifically (Table 2). Overall, most of the isolates resistant to ampicillin as determined using both disk diffusion (98%) and broth microdilution (94%) results harbored one of the beta-lactamase genes.

**TABLE 2.**
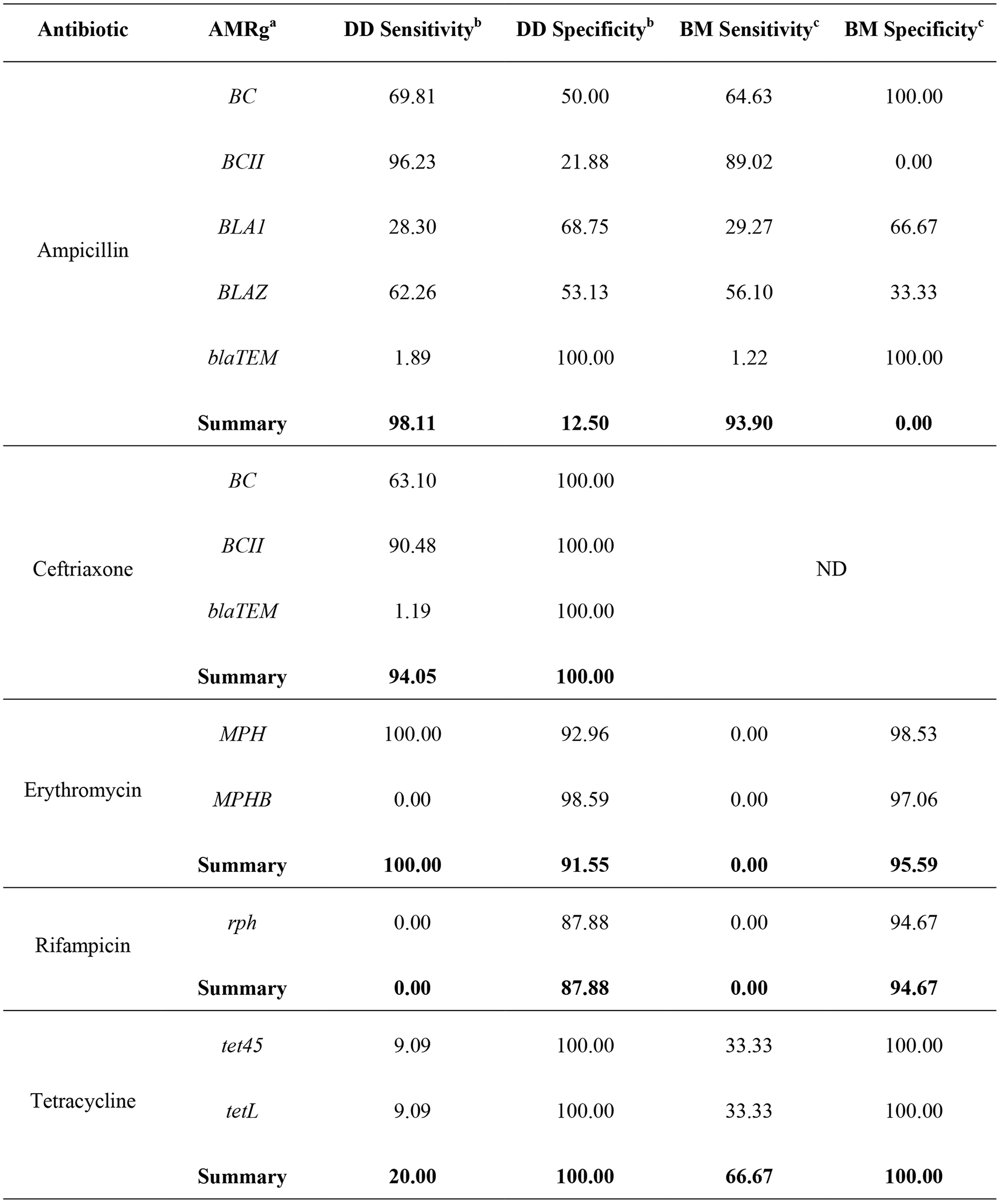
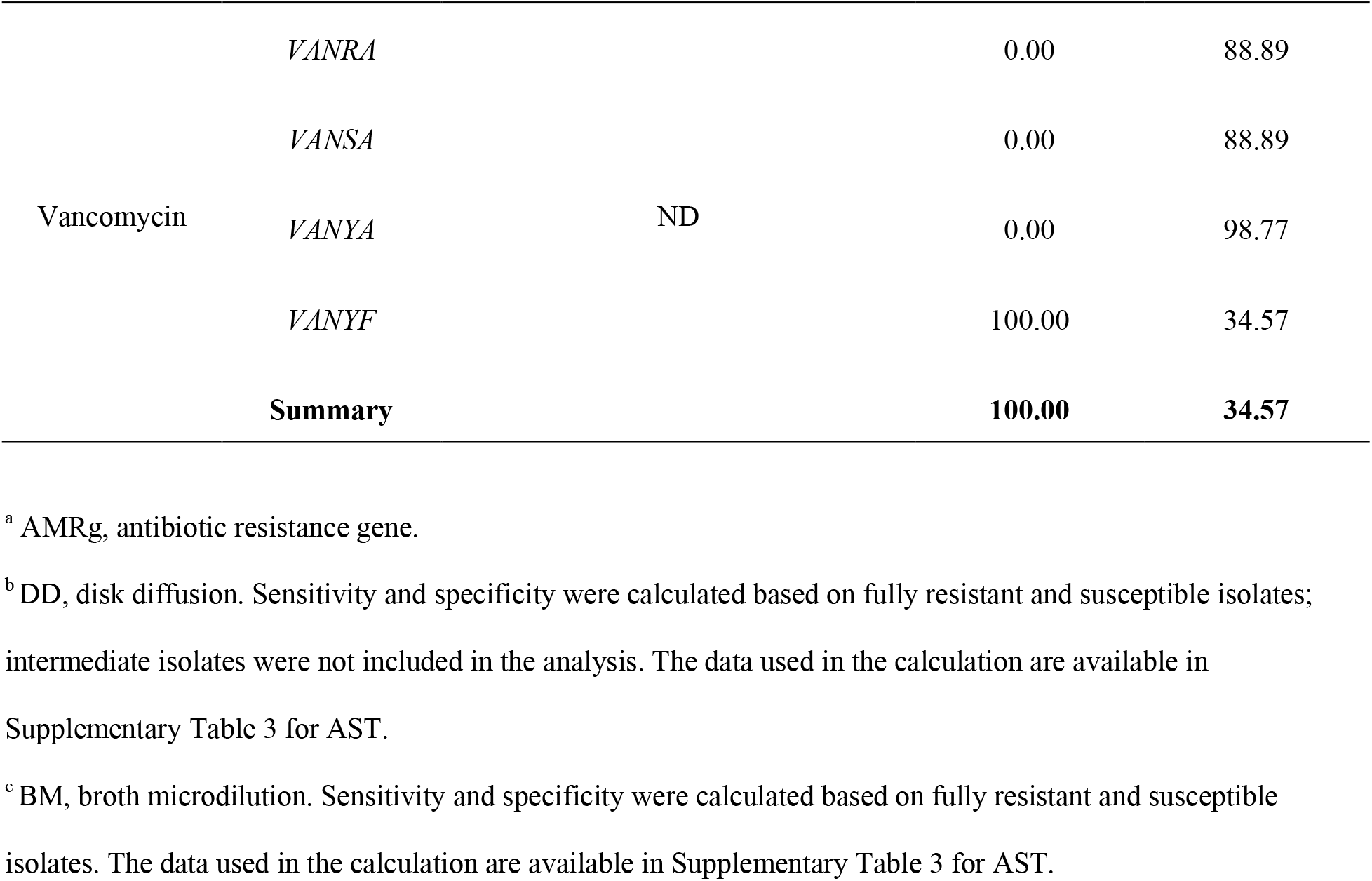
Sensitivity and specificity of antimicrobial resistance genes (AMRg) as markers of phenotypic resistance

Eleven isolates carried the *mph* (macrolide phosphotransferase) gene and two of these were also resistant to erythromycin as determined using the disk diffusion results, yielding 100% sensitivity (Table 2). However, the remaining five isolates that carried *mph* were susceptible to erythromycin, therefore producing a specificity of 92%. Based on the broth microdilution, no isolates were identified as resistant to erythromycin, resulting in a sensitivity of 0%. However, the specificity was high (96%), as nearly all isolates that carried *mph* gene (n = 10) were intermediately resistant. Based on both testing methods, *mphB* (n = 2) was found only in susceptible isolates, resulting in a sensitivity of 0% and specificity of 99% (disk diffusion) and 97% (broth microdilution).

Of the four isolates carrying the *rph* gene, which is known to confer resistance to rifampicin, none showed phenotypic resistance as determined with both disk diffusion and broth microdilution methods (Table 2). This resulted in 0% sensitivity, but high specificity (85% and 95%, respectively).

The sensitivity of *tet45* (n = 1) or *tetL* (n = 1) for predicting tetracycline resistance was low (9% for each gene) when using disk diffusion testing results (Table 2). The low sensitivity was due to only 2 out of 11 phenotypically resistant isolates (as determined using disk diffusion) carrying the tetracycline resistance genes. Interestingly, the sensitivity was substantially higher when broth microdilution results were used to interpret phenotypic resistance. Two of the three tetracycline-resistant isolates harbored either *tet45* or *tetL*, producing a combined sensitivity of 67%. Specificity was 100% based on the results of both disk diffusion and broth microdilution, as no susceptible isolates carried a *tet* gene.

In terms of vancomycin resistance, gene *vanYF* was detected in 57 isolates and was the only *van* gene detected in the three vancomycin-resistant isolates (Table 2). This resulted in a 100% sensitivity for the prediction of vancomycin resistance based on the presence of *vanYF* gene. However, *vanYF* was also detected in many vancomycin susceptible isolates, which resulted in a specificity of 34% (Table 2). Vancomycin resistance was not interpreted using disk diffusion results, hence we did not include it in this analysis.

Previous studies did not report associations between antimicrobial resistance gene presence and phenotypic resistance of isolates from the *B. cereus* group species. Few studies reported the presence of acquired antimicrobial resistance genes among studied isolates in combination with AST results (46, 47). Although the Bianco et al. study did not calculate the association between gene presence and phenotypic resistance, gene presence and phenotypic resistance were summarized (46). Bianco et al. identified beta-lactamases in all isolates and further characterized all isolates as phenotypically resistant to penicillin G and all isolates phenotypically non-susceptible to ceftriaxone (46). Although sensitivity and specificity were not reported in the Bianco et al. study, it is clear that the relationship between beta-lactamase presence and phenotypic resistance was strong, which is similar to our findings (46). Bianco et al. identified a single isolate that carried the *tetL* gene and this isolate was identified as intermediately resistant to tetracycline (no isolates were identified as resistant). In contrast, all of the *tet* positive isolates from our study were found to be phenotypically resistant isolates (46). The gene *mphB* was identified in 6 isolates by Bianco et al., but only 4 isolates were intermediately resistant to erythromycin showing *mphB* as a poor indicator of phenotypic resistance, similar to our findings (46). Lastly, *van* genes were identified in all isolates in the Bianco et al. study; however, all isolates were also phenotypically susceptible to vancomycin (46). Similarly, *van* genes were a poor predictor of vancomycin resistance for isolates tested in our study. Our findings, as well as Bianco et al. findings, show the need to further characterize the *B. cereus* group resistome and its effect on phenotypic resistance. Both of our studies only investigated acquired antimicrobial resistance genes and did not analyze point mutations and/or loss of function in housekeeping genes. Further study of the *Bacillus cereus* resistome is needed to better understand the underlying mechanisms of the *B. cereus* group isolates’ antimicrobial resistance.

## MATERIALS AND METHODS

### Bacterial cultures

A collection of 85 *B. cereus* group isolates that had been previously isolated from dairy foods and dairy-associated environments and whole-genome sequences were included in this study (10, 34, 35). A list of all isolates used in this study and their corresponding metadata is available in Supplemental Table S1. Isolates were resuscitated from cryo stocks preserved at - 80°C by streaking a loopful of each cryo stock onto brain heart infusion (BHI) agar (BD Difco) and incubating a streaked plate for 15-18 h at 32ºC. After completed incubation, a single isolated colony was sub-streaked onto a fresh BHI plate and incubated at the same conditions as outlined above. Subsequently, five isolated colonies were collected using a sterile loop and inoculated into 5 ml of BHI broth. Inoculated broths were incubated at 32°C for 15-18 hours (50). The culture adjusted to 1 - 2 × 10^8 was used for a disk diffusion assay described in the following section.

### Disk diffusion assay

Disk diffusion assay was performed on each isolate by following the Clinical and Laboratory Standards Institute (CLSI) guideline (50). A total of nine antimicrobials applied on sterile disks were tested: ampicillin (10 µg/disk), ceftriaxone (30 µg/disk), ciprofloxacin (5 µg/disk), erythromycin (15 µg/disk), gentamicin (10 µg/disk), rifampicin (5 µg/disk), tetracycline (30 µg/disk), trimethoprim and sulfamethoxazole (1.25 + 23.75 µg/disk), and vancomycin (30 µg/disk) (50). Disks were produced by applying an antibiotic onto a sterile disk to achieve the defined mass or were purchased pre-made (HardyDisk AST, Hardy Diagnostics). Ampicillin, erythromycin, gentamicin, and rifampicin (Hardy Diagnostics) antibiotics were applied onto disks in the lab by preparing stock concentrations and aseptically applying 20 µl of each antibiotic onto a disk to achieve the target mass of antibiotic per disk. Disks were first let dry in a biosafety cabinet for 20 minutes and were then stored at -20ºC until use. A detailed description of solvents and volumes used is listed in Supplemental Table 4. Ceftriaxone, ciprofloxacin, tetracycline, trimethoprim and sulfamethoxazole, and vancomycin antibiotics were purchased pre-applied on the disks and were stored at -20ºC until use.

Bacterial cultures prepared as described in the previous section were streaked in a lawn onto Muller Hinton (MH) agar (Becton Dickinson) using a sterile cotton swab (Medline Industries, Inc.) (50). Antimicrobial disks were placed onto the inoculated agar within 15 minutes of inoculation using sterile forceps (50). A maximum of three disks was applied on each individual agar plate to avoid overlaps in zones of inhibition that could compromise the accuracy of zone measurement. Inoculated plates with antibiotic disks were incubated at 35°C for 16-20 hours (50). *Bacillus cereus* type strain ATCC 14579 was included in each batch of tests for quality control. An uninoculated MH agar was also included in each batch of tests to control for medium sterility.

Inhibition zones were measured in millimeters using a ruler and interpreted using the guidelines and recommendations from CLSI and the European Committee on Antimicrobial Susceptibility Testing (EUCAST) (50, 51). EUCAST guidelines were followed in cases when it was difficult to interpret zones of inhibition due to unclear edges (51). Examples include cases when some colonies grew within the zone of inhibition, when double zones of inhibition are observed, and when the antibiotic trimethoprim-sulfamethoxazole produced atypical and challenging to interpret inhibition zones (51). Zones of inhibitions were interpreted using the CLSI M100-ED31:2021 *Staphylococcus* spp. breakpoints (50) for all antibiotics excluding ampicillin and ceftriaxone as they were undefined. Ampicillin and ceftriaxone results were interpreted using EUCAST v11.0 *Staphylococcus spp*. disk diffusion breakpoints (52). Vancomycin results were left interpreted as no breakpoints were defined in CLSI M100 or EUCAST v11.0 *Staphylococcus spp*.

### Broth microdilution assay

Broth microdilution assays were performed on each isolate following the M45-ED3 CLSI guideline (32). A total of 18 antimicrobials were tested: ampicillin (0.12-16 μg/mL), ceftriaxone (8-64 μg/mL), ciprofloxacin (0.5-2 μg/mL), clindamycin (0.12-2 μg/mL), daptomycin (0.25-8 μg/mL), erythromycin (0.25-4 μg/mL), gatifloxacin (1-8 μg/mL), gentamicin (2-500 μg/mL), levofloxacin (0.25-8 μg/mL), linezolid (0.5-8 μg/mL), oxacillin + 2% NaCl (0.25-8 μg/mL), penicillin (0.06-8 μg/mL), quinupristin & dalfopristin (0.12-4 μg/mL), rifampicin (0.5-4 μg/mL), streptomycin (1000 μg/mL), tetracycline (2-16 μg/mL), trimethoprim & sulfamethoxazole (0.5/9.5-4/76 μg/mL), and vancomycin (1-64 μg/mL). The Sensititre GPN3F 96-well plates pre-loaded with the appropriate antibiotics were used for broth microdilution testing (ThermoFisher). *Bacillus cereus* group isolates’ inocula were prepared from overnight cultures (see “Bacterial cultures” section) by adjusting their concentration to 1-2 × 10^8^ CFU/ml using a spectrophotometer (Eppendorf BioPhotometer 6131). A previously established OD-CFU standard curve was used to estimate the CFU/ml based on the OD reading. The inoculum was then dispensed in a GPN3F 96, 50 μL per well. Each plate included a positive control (no antibiotic added) and negative control (just MHB broth, no culture added). Inoculated plates were covered with a sealing tape provided with the Sensititre plates and incubated at 35ºC for 18-24 hours. To verify the inoculum concentration, dilutions of each culture suspension were spread plated onto BHI and incubated at 30ºC for 18-24 hours. The plates were counted to ensure that the concentration of each inoculum loaded into the Sensititre plates were between 10^5^ - 10^6^ CFU/ml.

Minimum inhibitory concentrations (MIC) were determined based on the guidelines and recommendations from CLSI (32). CLSI M07-A9 guideline was utilized in instances of unclear growth interpretations. This commonly included cases when a well had a very small amount of bacterial growth. In such cases, the bacterial pellets were measured in millimeters using a ruler and considered valid growth only if the pellet button was ≥ 2 mm (53). The MICs were interpreted using the M45-ED3 CLSI *Bacillus* spp. Breakpoints, excluding vancomycin which was interpreted using CLSI M100-ED31:2021 *Staphylococcus* spp. breakpoints (50).

### Genotyping, phylogenetic analysis, and detection of antimicrobial resistance genes using whole-genome sequencing

Sequencing reads for *Bacillus cereus* group isolates that had been previously sequenced were obtained from the NCBI SRA database (10, 34, 35). Sequence accession numbers are listed in Supplemental Table S1. The quality of sequences was examined with FASTQC v0.11.8 using default settings (54). Poor quality bases and adaptors were removed with Trimmomatic v0.39 using default settings and Nextera PE adapter sequences (55). Trimmed sequences were then assembled using SPAdes v3.15 with the kmer sizes of 99 and 127 and the careful option (56). Assembled genomes were checked for quality using QUAST v5.0.2 using the default parameters (57).

Draft genomes and processed reads were analyzed using Abricate v1.01 and Ariba v2.14.6, respectively, to detect the presence of acquired resistance genes (58, 59). MEGARes 2.0 and Resfinder databases were used with both programs. Gene presence was determined with each program using their respective default ≥ 90% percent coverage and identity.

BTyper3 v3.3.1 was used for taxonomic identification, genotyping (i.e., *panC, rpoB*, MLST typing) and identification of virulence genes using default settings (36). PROKKA v1.14.6 was used to annotate genomes and produce GFF files that were then used for the analysis of pangenome with Roary v3.11.2 (60, 61). The core genome alignment produced by Roary was used for phylogenetic inference with RAxML v8.2.12 (61, 62). RAxML was run with 1,000 bootstrap repetitions and the tree with the highest likelihood was used for visualization of phylogenetic relationships among isolates. The code for the bioinformatic analysis workflow outlined above is available at https://github.com/EmmaMills101/Bacillus-Cereus-Workflow.

### Data visualization and statistical analyses

Distributions of disk diffusion zones of inhibition and broth microdilution minimum inhibitory concentrations for each antibiotic were plotted in R and R Studio v4.0.4 using packages base v4.0.4, graphics v4.0.4, ggplot v3.3.3, grDevices v4.0.4, methods v4.0.4, and stats v4.0.4 (63). Differences in the resistant prevalence between the two methods were calculated using the difference in proportion hypothesis testing with 0.05 significance in R (hypothesestest v1.0 package) (63). Phylogenetic tree, sequence types, virulence genes, antibiotic susceptibility testing results, and antibiotic resistance genetic determinants were visualized using iTOL v5 (64). The sensitivity and specificity for predicting phenotypic antibiotic resistance based on detected antimicrobial resistance genes were calculated using the guidelines outlined in Trevethan, 2017 (65).

## ACKNOWLEDGMENTS

This work was supported by the Penn State College of Agricultural Science Undergraduate research grants awarded to Emma Mills and Erin Sullivan, and by the USDA National Institute of Food and Agriculture Hatch Appropriations under Project #PEN04646 and Accession #1015787.

## REFERENCES

1. Carroll LM, Wiedmann M, Mukherjee M, Nicholas DC, Mingle LA, Dumas NB, Cole JA, Kovac J. 2019. Characterization of Emetic and Diarrheal Bacillus cereus Strains From a 2016 Foodborne Outbreak Using Whole-Genome Sequencing: Addressing the Microbiological, Epidemiological, and Bioinformatic Challenges. Front Microbiol 10.

2. Owusu-Kwarteng J, Wuni A, Akabanda F, Tano-Debrah K, Jespersen L. 2017. Prevalence, virulence factor genes and antibiotic resistance of Bacillus cereus sensu lato isolated from dairy farms and traditional dairy products. BMC Microbiol 17:65.

3. Scallan E, Hoekstra RM, Angulo FJ, Tauxe RV, Widdowson M-A, Roy SL, Jones JL, Griffin PM. 2011. Foodborne illness acquired in the United States--major pathogens. Emerg Infect Dis 17:7–15.

4. Kim Y-J, Kim H-S, Kim K-Y, Chon J-W, Kim D-H, Seo K-H. 2016. High Occurrence Rate and Contamination Level of Bacillus cereus in Organic Vegetables on Sale in Retail Markets. Foodborne Pathog Dis 13:656–660.

5. Kotzekidou P. 2013. Microbiological examination of ready-to-eat foods and ready-to-bake frozen pastries from university canteens. Food Microbiol 34:337–343.

6. Carroll LM, Wiedmann M, Kovac J. Proposal of a Taxonomic Nomenclature for the Bacillus cereus Group Which Reconciles Genomic Definitions of Bacterial Species with Clinical and Industrial Phenotypes. mBio 11:e00034–20.

7. Carroll LM, Cheng, RA, Kovac J. No Assembly Required: Using BTyper3 to Assess the Congruency of a Proposed Taxonomic Framework for the Bacillus cereus Group with Historical Typing Methods. Frontiers in Microbiology, https://doi.org/10.3389/fmicb.2020.580691.

8. Miller RA, Jian J, Beno SM, Wiedmann M, Kovac J. 2018. Intraclade Variability in Toxin Production and Cytotoxicity of Bacillus cereus Group Type Strains and Dairy-Associated Isolates. Appl Environ Microbiol 84:e02479–17.

9. Guinebretière M-H, Velge P, Couvert O, Carlin F, Debuyser M-L, Nguyen-The C. 2010. Ability of Bacillus cereus Group Strains to Cause Food Poisoning Varies According to Phylogenetic Affiliation (Groups I to VII) Rather than Species Affiliation. J Clin Microbiol 48:3388–3391.

10. Miller RA, Jian J, Beno SM, Wiedmann M, Kovac J. 2018. Intraclade Variability in Toxin Production and Cytotoxicity of Bacillus cereus Group Type Strains and Dairy-Associated Isolates. Appl Environ Microbiol 84:e02479–17.

11. Scallan E, Hoekstra RM, Angulo FJ, Tauxe RV, Widdowson M-A, Roy SL, Jones JL, Griffin PM. 2011. Foodborne Illness Acquired in the United States—Major Pathogens. Emerg Infect Dis 17:7–15.

12. Dierick K, Van Coillie E, Swiecicka I, Meyfroidt G, Devlieger H, Meulemans A, Hoedemaekers G, Fourie L, Heyndrickx M, Mahillon J. 2005. Fatal Family Outbreak of Bacillus cereus-Associated Food Poisoning. J Clin Microbiol 43:4277–4279.

13. Naranjo M, Denayer S, Botteldoorn N, Delbrassinne L, Veys J, Waegenaere J, Sirtaine N, Driesen RB, Sipido KR, Mahillon J, Dierick K. 2011. Sudden Death of a Young Adult Associated with Bacillus cereus Food Poisoning. J Clin Microbiol 49:4379–4381.

14. Colaco CMG, Basile K, Draper J, Ferguson PE. 2021. Fulminant Bacillus cereus food poisoning with fatal multi-organ failure. BMJ Case Rep CP 14:e238716.

15. Glasset B, Herbin S, Granier SA, Cavalié L, Lafeuille E, Guérin C, Ruimy R, Casagrande-Magne F, Levast M, Chautemps N, Decousser J-W, Belotti L, Pelloux I, Robert J, Brisabois A, Ramarao N. 2018. Bacillus cereus, a serious cause of nosocomial infections: Epidemiologic and genetic survey. PLoS ONE 13:e0194346.

16. Veysseyre F, Fourcade C, Lavigne J-P, Sotto A. 2015. Bacillus cereus infection: 57 case patients and a literature review. Med Mal Infect 45:436–440.

17. Yamada K, Shigemi H, Suzuki K, Yasutomi M, Iwasaki H, Ohshima Y. 2019. Successful management of a Bacillus cereus catheter-related bloodstream infection outbreak in the pediatric ward of our facility. J Infect Chemother Off J Jpn Soc Chemother 25:873–879.

18. Saikia L, Gogoi N, Das PP, Sarmah A, Punam K, Mahanta B, Bora S, Bora R. 2019. Bacillus cereus-Attributable Primary Cutaneous Anthrax-Like Infection in Newborn Infants, India. Emerg Infect Dis 25:1261–1270.

19. Gargis AS, McLaughlin HP, Conley AB, Lascols C, Michel PA, Gee JE, Marston CK, Kolton CB, Rodriguez-R LM, Hoffmaster AR, Weigel LM, Sue D. Analysis of Whole-Genome Sequences for the Prediction of Penicillin Resistance and β-Lactamase Activity in Bacillus anthracis. mSystems 3:e00154–18.

20. Aygun FD, Aygun F, Cam H. 2016. Successful Treatment of Bacillus cereus Bacteremia in a Patient with Propionic Acidemia. Case Rep Pediatr 2016:6380929.

21. Ikeda M, Yagihara Y, Tatsuno K, Okazaki M, Okugawa S, Moriya K. 2015. Clinical characteristics and antimicrobial susceptibility of Bacillus cereus blood stream infections. Ann Clin Microbiol Antimicrob 14.

22. Schaefer G, Campbell W, Jenks J, Beesley C, Katsivas T, Hoffmaster AR, Mehta SR, Reed S. Persistent Bacillus cereus Bacteremia in 3 Persons Who Inject Drugs, San Diego, California, USA - Volume 22, Number 9—September 2016 - Emerging Infectious Diseases journal - CDC https://doi.org/10.3201/eid2209.150647.

23. Zhang Y, Chen M, Yu P, Yu S, Wang J, Guo H, Zhang J, Zhou H, Chen M, Zeng H, Wu S, Pang R, Ye Q, Xue L, Zhang S, Li Y, Zhang J, Wu Q, Ding Y. 2020. Prevalence, Virulence Feature, Antibiotic Resistance and MLST Typing of Bacillus cereus Isolated from Retail Aquatic Products in China. Front Microbiol 0.

24. Materon IC, Queenan AM, Koehler TM, Bush K, Palzkill T. 2003. Biochemical Characterization of β-Lactamases Bla1 and Bla2 from Bacillus anthracis. Antimicrob Agents Chemother 47:2040–2042.

25. Karsisiotis AI, Damblon CF, Roberts GCK. 2013. Solution structures of the Bacillus cereus metallo-β-lactamase BcII and its complex with the broad spectrum inhibitor R-thiomandelic acid. Biochem J 456:397–407.

26. Savić D, Miljković-Selimović B, Lepšanović Z, Tambur Z, Konstantinović S, Stanković N, Ristanović E. 2016. Antimicrobial susceptibility and β-lactamase production in Bacillus cereus isolates from stool of patients, food and environment samples. Vojnosanit Pregl 73:904–909.

27. Shimoyama Y, Umegaki O, Ooi Y, Agui T, Kadono N, Minami T. 2017. Bacillus cereus pneumonia in an immunocompetent patient: a case report. JA Clin Rep 3:25.

28. Katsuya H, Takata T, Ishikawa T, Sasaki H, Ishitsuka K, Takamatsu Y, Tamura K. 2009. A patient with acute myeloid leukemia who developed fatal pneumonia caused by carbapenem-resistant Bacillus cereus. J Infect Chemother Off J Jpn Soc Chemother 15:39–41.

29. Shawish R, Tarabees R. 2017. Prevalence and antimicrobial resistance of Bacillus cereus isolated from beef products in Egypt. Open Vet J 7:337–341.

30. Fiedler G, Schneider C, Igbinosa EO, Kabisch J, Brinks E, Becker B, Stoll DA, Cho G-S, Huch M, Franz CMAP. 2019. Antibiotics resistance and toxin profiles of Bacillus cereus-group isolates from fresh vegetables from German retail markets. BMC Microbiol 19:250.

31. Zhao S, Tyson GH, Chen Y, Li C, Mukherjee S, Young S, Lam C, Folster JP, Whichard JM, McDermott PF. 2016. Whole-Genome Sequencing Analysis Accurately Predicts Antimicrobial Resistance Phenotypes in Campylobacter spp. Appl Environ Microbiol 82:459–466.

32. Carroll LM, Wiedmann M, den Bakker H, Siler J, Warchocki S, Kent D, Lyalina S, Davis M, Sischo W, Besser T, Warnick LD, Pereira RV. 2017. Whole-Genome Sequencing of Drug-Resistant Salmonella enterica Isolates from Dairy Cattle and Humans in New York and Washington States Reveals Source and Geographic Associations. Appl Environ Microbiol 83:e00140–17.

33. Frenzel E, Kranzler M, Stark TD, Hofmann T, Ehling-Schulz M. 2015. The Endospore-Forming Pathogen Bacillus cereus Exploits a Small Colony Variant-Based Diversification Strategy in Response to Aminoglycoside Exposure. mBio 6:e01172–15.

34. Kovac J, Miller RA, Carroll LM, Kent DJ, Jian J, Beno SM, Wiedmann M. 2016. Production of hemolysin BL by Bacillus cereus group isolates of dairy origin is associated with whole-genome phylogenetic clade. BMC Genomics 17:581.

35. Miller RA, Beno SM, Kent DJ, Carroll LM, Martin NH, Boor KJ, Kovac J. 2016. Bacillus wiedmannii sp. nov., a psychrotolerant and cytotoxic Bacillus cereus group species isolated from dairy foods and dairy environments. Int J Syst Evol Microbiol 66:4744–4753.

36. Carroll LM, Kovac J, Miller RA, Wiedmann M. 2017. Rapid, High-Throughput Identification of Anthrax-Causing and Emetic Bacillus cereus Group Genome Assemblies via BTyper, a Computational Tool for Virulence-Based Classification of Bacillus cereus Group Isolates by Using Nucleotide Sequencing Data. Appl Environ Microbiol 83.

37. Yu S, Yu P, Wang J, Li C, Guo H, Liu C, Kong L, Yu L, Wu S, Lei T, Chen M, Zeng H, Pang R, Zhang Y, Wei X, Zhang J, Wu Q, Ding Y. 2020. A Study on Prevalence and Characterization of Bacillus cereus in Ready-to-Eat Foods in China. Front Microbiol 10:3043.

38. Premkrishnan BNV, Heinle CE, Uchida A, Purbojati RW, Kushwaha KK, Putra A, Santhi PS, Khoo BWY, Wong A, Vettath VK, Drautz-Moses DI, Junqueira ACM, Schuster SC. 2021. The genomic characterisation and comparison of Bacillus cereus strains isolated from indoor air. Gut Pathog 13:6.

39. The Comprehensive Antibiotic Resistance Database. https://card.mcmaster.ca, Accessed on 11/20/2021.

40. Rasia RM, Vila AJ. 2004. Structural Determinants of Substrate Binding to Bacillus cereus Metallo-β-lactamase*. J Biol Chem 279:26046–26051.

41. Qi X, Lin W, Ma M, Wang C, He Y, He N, Gao J, Zhou H, Xiao Y, Wang Y, Zhang P. 2016. Structural basis of rifampin inactivation by rifampin phosphotransferase. Proc Natl Acad Sci 113:3803–3808.

42. Stogios PJ, Cox G, Spanogiannopoulos P, Pillon MC, Waglechner N, Skarina T, Koteva K, Guarné A, Savchenko A, Wright GD. 2016. Rifampin phosphotransferase is an unusual antibiotic resistance kinase. Nat Commun 7:11343.

43. Thompson MK, Keithly ME, Harp J, Cook PD, Jagessar KL, Sulikowski GA, Armstrong RN. 2013. Structural and chemical aspects of resistance to the antibiotic fosfomycin conferred by FosB from Bacillus cereus. Biochemistry 52:7350–7362.

44. Read TD, Peterson SN, Tourasse N, Baillie LW, Paulsen IT, Nelson KE, Tettelin H, Fouts DE, Eisen JA, Gill SR, Holtzapple EK, Økstad OA, Helgason E, Rilstone J, Wu M, Kolonay JF, Beanan MJ, Dodson RJ, Brinkac LM, Gwinn M, DeBoy RT, Madpu R, Daugherty SC, Durkin AS, Haft DH, Nelson WC, Peterson JD, Pop M, Khouri HM, Radune D, Benton JL, Mahamoud Y, Jiang L, Hance IR, Weidman JF, Berry KJ, Plaut RD, Wolf AM, Watkins KL, Nierman WC, Hazen A, Cline R, Redmond C, Thwaite JE, White O, Salzberg SL, Thomason B, Friedlander AM, Koehler TM, Hanna PC, Kolstø A-B, Fraser CM. 2003. The genome sequence of Bacillus anthracis Ames and comparison to closely related bacteria. Nature 423:81–86.

45. Ligozzi M, Lo Cascio G, Fontana R. 1998. vanA Gene Cluster in a Vancomycin-Resistant Clinical Isolate of Bacillus circulans. Antimicrob Agents Chemother 42:2055–2059.

46. Bianco A, Capozzi L, Monno MR, Del Sambro L, Manzulli V, Pesole G, Loconsole D, Parisi A. 2021. Characterization of Bacillus cereus Group Isolates from Human Bacteremia by Whole-Genome Sequencing. Front Microbiol 11:3273.

47. Zhu K, Hölzel CS, Cui Y, Mayer R, Wang Y, Dietrich R, Didier A, Bassitta R, Märtlbauer E, Ding S. 2016. Probiotic Bacillus cereus Strains, a Potential Risk for Public Health in China. Front Microbiol 7:718.

48. Park Y-B, Kim J-B, Shin S-W, Kim J-C, Cho S-H, Lee B-K, Ahn J, Kim J-M, Oh D-H. 2009. Prevalence, genetic diversity, and antibiotic susceptibility of Bacillus cereus strains isolated from rice and cereals collected in Korea. J Food Prot 72:612–617.

49. Yu P, Yu S, Wang J, Guo H, Zhang Y, Liao X, Zhang J, Wu S, Gu Q, Xue L, Zeng H, Pang R, Lei T, Zhang J, Wu Q, Ding Y. 2019. Bacillus cereus Isolated from Vegetables in China: Incidence, Genetic Diversity, Virulence Genes, and Antimicrobial Resistance. Front Microbiol 10:948.

50. Weinstein M, Lewis J, Bobenchick A, Campeau S, Cullen S, Galas M, Gold H, Humphries R, Kirn T, Limbago B, Mathers A, Mazzulli T, Richter S, Satlin M, Schuetz A, Simner P. 2021. CLSI M100 ED31:2021 Performance Standards for Antimicrobial Susceptibility Testing, 31st Edition. Accessed 9 July 2021.

51. 2021. EUCAST: Disk diffusion methodology. Disk Diffus - Read Guide V 80. The Eurepean Committee on Antimicrobial Susceptibility Testing.

52. 2021. EUCAST: Clinical breakpoints and dosing of antibiotics. Eur Comm Antimicrob Susceptibility Test - EUCAST 2021. Accessed 9 July 2021.

53. Adler J, Cockerill F, Dudley M, Eliopoulos G, Hardy D, Hecht D, Hindler J, Patel J, Powell M, Swenson J, Thomson R, Traczewksi M, Turnidge J, Weinstein M, Wikler M, Zimmer B. CLSI M07 ED09:2012 Methods for Dilution Antimicrobial Susceptibility Tests for Bacteria That Grow Aerobically; Approved Standard, Ninth Edition 32.

54. Babraham Bioinformatics - FastQC A Quality Control tool for High Throughput Sequence Data. https://www.bioinformatics.babraham.ac.uk/projects/fastqc/, Accessed on 11/20/2021.

55. Bolger AM, Lohse M, Usadel B. 2014. Trimmomatic: a flexible trimmer for Illumina sequence data. Bioinformatics 30:2114–2120.

56. Bankevich A, Nurk S, Antipov D, Gurevich AA, Dvorkin M, Kulikov AS, Lesin VM, Nikolenko SI, Pham S, Prjibelski AD, Pyshkin AV, Sirotkin AV, Vyahhi N, Tesler G, Alekseyev MA, Pevzner PA. 2012. SPAdes: a new genome assembly algorithm and its applications to single-cell sequencing. J Comput Biol J Comput Mol Cell Biol 19:455–477.

57. Gurevich A, Saveliev V, Vyahhi N, Tesler G. 2013. QUAST: quality assessment tool for genome assemblies. Bioinformatics 29:1072–1075.

58. Seemann T. 2021. tseemann/abricate. Perl.

59. Hunt M, Mather AE, Sánchez-Busó L, Page AJ, Parkhill J, Keane JA, Harris SR. 2017. ARIBA: rapid antimicrobial resistance genotyping directly from sequencing reads. Microb Genomics 3.

60. Seemann T. 2014. Prokka: rapid prokaryotic genome annotation. Bioinforma Oxf Engl 30:2068–2069.

61. Page AJ, Cummins CA, Hunt M, Wong VK, Reuter S, Holden MTG, Fookes M, Falush D, Keane JA, Parkhill J. 2015. Roary: rapid large-scale prokaryote pan genome analysis. Bioinforma Oxf Engl 31:3691–3693.

62. Stamatakis A. 2014. RAxML version 8: a tool for phylogenetic analysis and post-analysis of large phylogenies. Bioinformatics 30:1312–1313.

63. 2015. RStudio: Integrated Development for R. RStudio. RStudio.

64. Letunic I, Bork P. 2016. Interactive tree of life (iTOL) v3: an online tool for the display and annotation of phylogenetic and other trees. Nucleic Acids Res 44:W242–245.

65. Trevethan R. 2017. Sensitivity, Specificity, and Predictive Values: Foundations, Pliabilities, and Pitfalls in Research and Practice. Front Public Health 5.

